# Generative AI impact on protein stability prediction in breast cancer genes

**DOI:** 10.1101/2024.06.03.597089

**Authors:** Rohan Gnanaolivu, Steven N Hart

## Abstract

The functional classification of a missense variant in cancer predisposition genes is often challenging due to how rare the variant is observed in the population. When available, clinicians utilize a combination of family history, *in vitro* functional assays and *in silico* methods to infer protein function. *In silico* methods, such as missense predictors (predict changes in protein function) and protein stability predictors (predict changes in free energy) have been used to help classify a missense variant in accordance with the American College of Medical Genetics and Genomics (ACMG) guideline. To measure protein stability, many *in silico* algorithms predict stability based on the change of free energy and most accurate protein stability predictors require a wild-type protein template. In this study, we examine the use of generative AI to predict high-resolution protein structures as templates analyzed with protein stability methods to evaluate loss of function (LOF) activity in cancer predisposition genes *BRCA1, BRCA2, PALB2* and *RAD51C* upon the presence of missense variant. Utilizing multiplexed assay of variant effect measurements and variant classifications from ClinVar, we find that prediction of Gibbs free energy (ΔΔG) from AlphaFold2 (AF2) structures analyzed with FoldX predicts LOF better than experimental-derived wild type structures in the BRCT domain of *BRCA1* and the DNA binding domain (DBD) of *BRCA2*, but not in *PALB2* and *RAD51C*. We also find that AF2 structures in the BRCT domain of BRCA1 and DBD-Dss1 domain of BRCA2 analyzed with FoldX measure homologous DNA recombination (HDR) activity significantly better than Rosetta and DDGun3D. Our study also revealed that there are other factors that contribute to predicting loss of function activity other than protein stability, with AlphaMissense ranking the best overall predictor of LOF activity in these tumor suppressor breast cancer genes.

**Author Summary:** The stability of a protein, often expressed in terms of Gibbs free energy (ΔΔG), is a critical factor in predicting loss of function (LOF) activity when a missense variant is present. The effect is higher in haploinsufficient genes like the tumor suppressor genes *BRCA1, BRCA2, PALB2* and *RAD51C*. Protein stability predictors that utilizes a wild-type structure to make its predictions is often limited by the availability of experimentally-derived protein structures. Here, in our study we show that generative AI, like AlphaFold2 (AF2) can predict structures similar to experimentally-derived structures with high similarity. Furthermore, protein stability tools such as FoldX, Rosetta, and DDGun3D can be used in conjunction to measure changes in stability. From our study, we find that complex AF2 structures representing the BRCT domain of *BRCA1* and DBD domain of *BRCA2* analyzed by FoldX predicts function significantly better than the experimentally-derived structures. However, predicted |ΔΔG| does not predict function better than purpose-built *in silico* missense predictors for protein function. Overall, we find the AlphaMissense is the best predictor to predict function in these tumor suppressor breast cancer genes.

## Introduction

Tumor suppressor genes *BRCA1, BRCA2, PALB2*, and *RAD51C* play crucial roles in homology-directed repair (HDR) activity, and mutations in these genes have been implicated in breast cancer[1]. While the functional impact of truncating mutations in these genes are well characterized, the clinical impact of >95% of all possible missense mutation in genes *BRCA2, PALB2* and *RAD51C* and > 80% of all possible missense mutations in *BRCA1* remain unclassified, therefore, most are commonly referred to as variants of uncertain significance[2]. Various *in silico* and *in vitro* functional assays have been employed to evaluate function. Commonly used *in silico* methods primarily use features such as sequence conversation, protein conformation and stereochemical properties of the amino acid as features to infer functional outcomes[3-5]. Researchers and Clinician have also utilized *in silico* stability predictors, which predict the destabilization or over-stabilization of the protein to infer loss or gain of function. Consequently, the absolute value of predicted ΔΔG is used to evaluate protein function [6-8].

Calculating protein stability involves measuring the effect and change in free energy of the protein based on the presence of a mutation. The change in protein structure directly impacts overall stability, particularly in haploinsufficient genes like *BRCA1, BRCA2, PALB2*, and *RAD51C*[9-12]. One method to assess protein stability is by estimating the difference in free energy of unfolding of the proteins (ΔG) between the wild-type and variant protein: ΔΔG = ΔG_variant – ΔG_wild-type. FoldX[13], Rosetta[14], and DDGun3D[15] are well-known stability predictors that induce a mutation from a given wild-type protein structure and predict ΔΔG. Recent publications have demonstrated that FoldX, Rosetta and DDGun3D had the highest correlation with functional measurements from deep mutational scanning data utilizing experimentally-derived protein complexes from the protein data bank (PDB) compared to other stability predictors[16]. However, the impact of stability on function is protein dependent, as there are several factors such as sequence composition, post-translation modification, haploinsufficiency, and binding partners that influence stability of the protein[17, 18]. Therefore, the predictive performance of these stability methods to predict LOF in these tumor suppressor breast cancer genes is still relatively unknown. The study of protein function using predicted protein stability involves several limitations as well, such as the inherent variability of these methods [19] and the availability of experimentally-derived crystalized structures of the complete protein.

Advances in generative AI, particularly in protein prediction models like AF2[20] and ESMFold [21], have shown promise in predicting structures that are highly similar to structures found in the PDB. AF2 and ESMFold were shown to predict protein structures that are highly similar to experimentally-derived structures in the Critical Assessment of Structure Prediction (CASP)15 challenge[22]. Recent CASP13[23] and CASP14[24] challenges highlighted the giant leap in advancement in protein structure prediction, with results highlighting the accuracy of predicted protein structures. Recent studies have demonstrated that features extracted from AF2 structures are effective in predicting the functional classification of missense variants [25]. However, the ability of using generative AI in HDR pathway genes to predict protein stability is relatively unknown and the impact of features derived from these generative AI models on protein stability is not well understood.

Experimental methods, such as multiplexed assay of variant effect (MAVE) and functional assays provide insight on LOF activity of *BRCA1, BRCA2, RAD51C*, and *PALB2*[26-31]. MAVE assays in these genes measure the HDR activity based on survival or growth of the cells upon the induced mutations. Results from these assays provide a reliable set of mutations to evaluate protein stability on the functional activity in our genes of interest [32]. However, these MAVE assays are shown to have high stochasticity and hence multiple replicates are often required. MAVE assays are also limited based on the biological mechanism involved and do not provide mutational classification on all possible missense mutations in these genes, with some assays being domain specific to the gene of interest.

*In silico* missense predictors are considered the weakest form of evidence to classify a missense variant[5]. REVEL [33] and BayesDel [34] are two *in silico* missense prediction tools that have been historically cited to accurately predict deleteriousness [35]. Newer, deep learning models such as AlphaMissense [36] and MetaRNN [37] have gained attention in predicting LOF activity. The effectiveness of these newer predictors, as well as those predicting protein stability based on functional classifications from MAVE experiments in these genes, is still largely unexplored.

The goal of this study was to assess how protein stability affects protein function in HDR genes. We used predicted protein structures from generative AI tools as the baseline wild-type structural templates, which were analyzed with FoldX, Rosetta, and DDGun3D to evaluate loss of function (LOF) activity in the genes *BRCA1, BRCA2, PALB2*, and *RAD51C*. Our results demonstrate that FoldX, Rosetta, and DDGun3D can predict stability using AF2 structures similar to experimentally-derived structures found in the PDB. Furthermore, AF2 wild-type structures enhance LOF predictions analyzed with FoldX in the BRCT domain of *BRCA1* and the DBD domain in BRCA2, compared with the experimentally-derived structures. FoldX was also shown to significantly outperform Rosetta and DDGun3D to predict HDR activity from the MAVE assays in the BRCT domain of *BRCA1* and the DBD domain in *BRCA2*. Finally, *in silico* predictor AlphaMissense ranked as the top predictor to predict LOF activity based on the Area Under the Curve (AUC) in the ordered regions of *BRCA1* and *BRCA2*, as well as it was found to be one of the top predictors in *PALB2* and *RAD51C*.

## Results

### Comparison of Generative AI protein prediction structures

From the literature, two protein structures are considered similar if the Root Mean Square Deviation (RMSD) between carbon-alpha (Cα) back-bone chains that are superimposed is <3.8Å [38]. Using this threshold as the metric for similarity, AF2 generates 3D protein structures comparatively similar to the experimentally-derived structures compared to ESMFold. A RMSD of <3.8Å threshold was generated for only 3 structures (4OFB[39], 1JNX[40] and 2W18[41]) by ESMFold, and due to the size limitation of 1024 residues, 2 complex structures (7LYB[42], 1IYJ[43]) were not analyzed. AF2 predicts 7 structures less than the RMSD threshold and only one 1IYJ fails the threshold (Table 1). 1IYJ could not be predicted without significant error and hence it was excluded from further analysis. Calculating the per residue distance (Å) when the predicted structure was superimposed on the experimentally-derived structure, >98% of the residues was <3.8Å in 6 out of 8 structures, highlighting the prediction accuracy of AF2. However, 7LYB had a mean residue distance of 19Å, even though the distances between the *BRCA1* Cα back-bone chain and its experimentally-derived counterpart was <3.8Å.

**Table 1:**
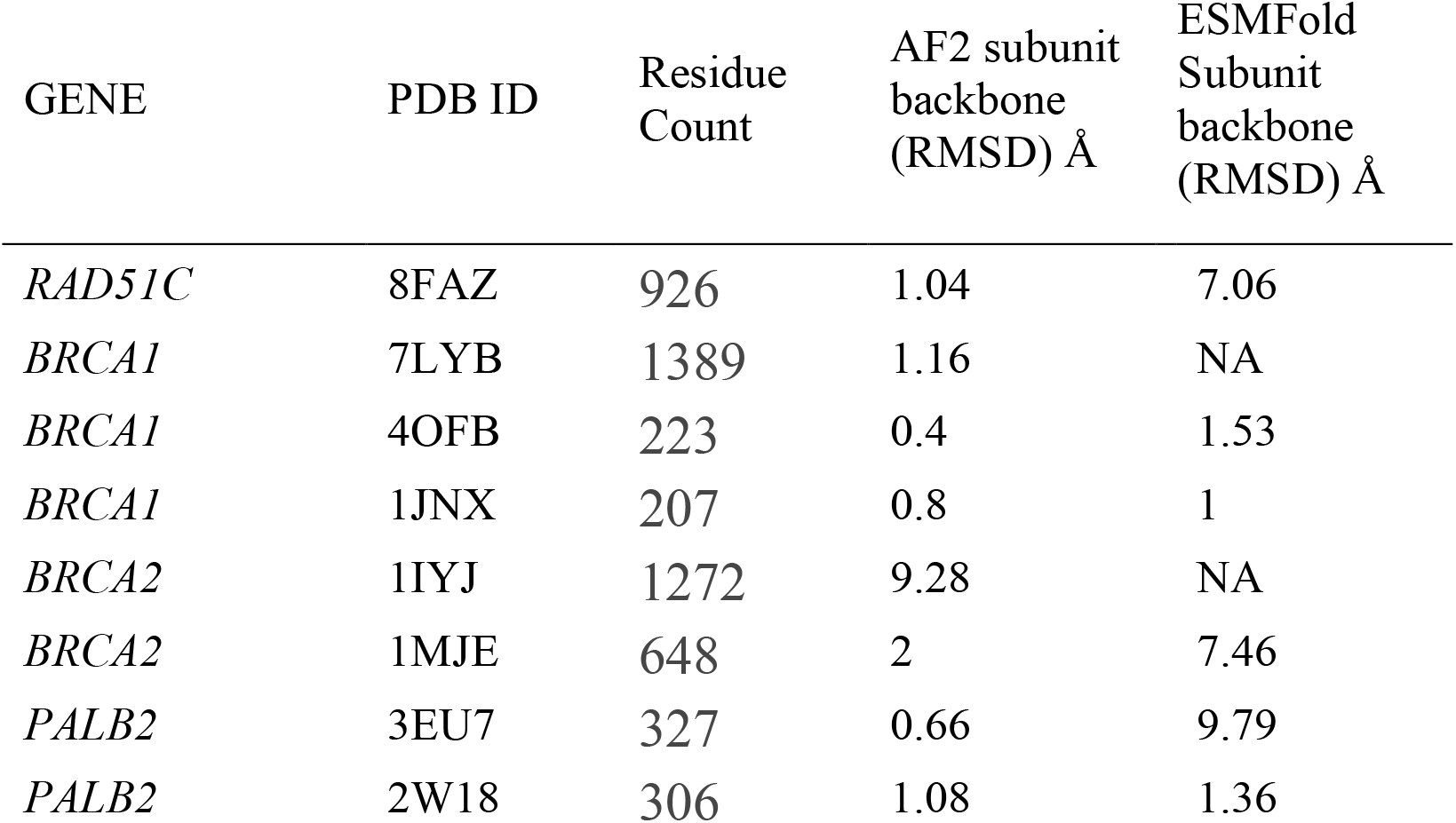
Root Mean Square Deviation (RMSD) values of AlphaFold2 and ESMFold structures superimposed on experimentally-derived structures from the PDB.

### Association of protein function from predicted |ΔΔG| generated from experimentally derived structures

Utilizing the Mann-Whitney U statistical test, the association of |ΔΔG| derived from 7 experimentally derived structures PDB ID:1JNX, 4OFB, 1JME, 3EU7[41], 2W18, and 8FAZ[44] representing regions of *BRCA1, BRCA2, PALB2* and *RAD51C*, analyzed with protein stability predictors FoldX, Rosetta and DDGun3D, was tested against the LOF classification from the mutations found in the ClinVar database and the MAVE assays. Our results show that there was a significant association of predicted |ΔΔG| to LOF activity in all 4 genes (S1 Fig).

### Linearity of |ΔΔG| generated from protein stability predictions using AF2 structures compared to experimentally-derived structures

Utilizing the 7 AF2 structures that had an RMSD of the Cα backbone < 3.8Å, the spearman correlation coefficient (rho) was calculated from the |ΔΔG| analyzed with the AF2 wild-type structures and experimentally-derived structures. There is a strong correlation between the |ΔΔG| predictions from FoldX, Rosetta, and DDGun3D using both AF2 structures and experimentally-derived structures (Fig. 1). FoldX and Rosetta had a rho value > 0.75 in 5 out of the 7 structures, indicating strong correlations, with 7LYB and IMJE being the only exception. FoldX generated a rho value of 0.63 (95% Confidence Interval (CI) 0.60-0.66) with 7LYB and 0.70 (95% CI 0.69-0.70) with 1MJE. Rosetta had a rho value of 0.71 (95% CI 0.68-0.73) with 7LYB and 0.75 with 1MJE (95% CI: 0.74-0.75). DDGun3D showed a strong correlation for all 7 structures, with a mean rho value >0.97.

**Fig 1:**
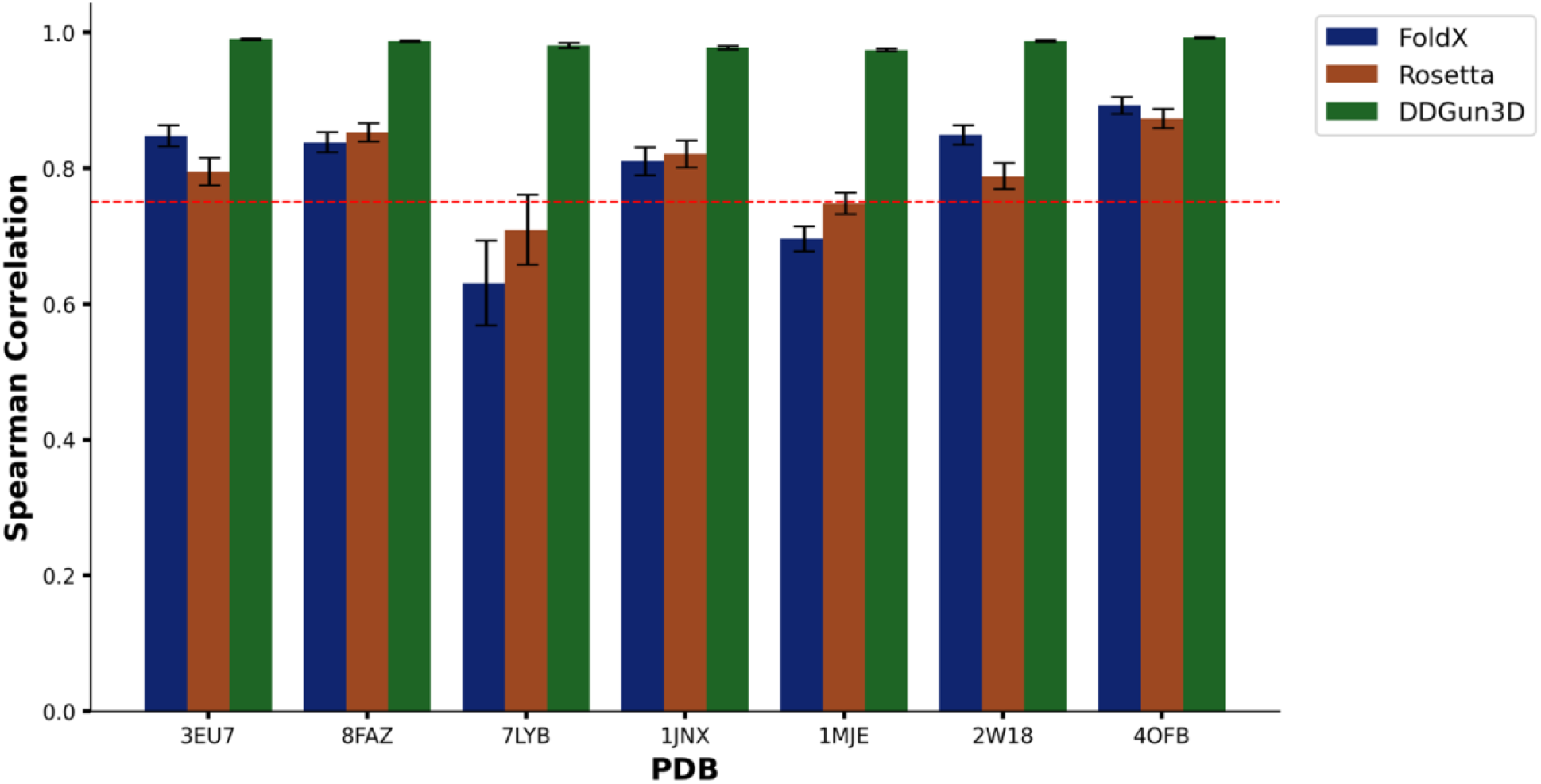
Spearman Correlation between predicted |ΔΔG| from FoldX, Rosetta and DDGun3D using experimentally-derived structurea vs AF2 structures as the wild-type template. The red line at y=0.75 indicates the threshold for strong correlation.

To test if the heterogeneity of the rho value was impacted by the features from AF2 structures, the linearity of the |ΔΔG| deltas calculated from predicted |ΔΔG| derived from AF2 structures and experimentally-derived structures analyzed with FoldX, Rosetta and DDGun3D with the features from AF2 structures was tested to better understand the impact of these features on the change of stability. A rho value was calculated, and we found no linearity on the deltas from the |ΔΔG| predictions with the AF2 features (S2A Fig). To understand if the heterogeneity of the |ΔΔG| deltas was dependent on the per residue distance in Å of the superimposed AF2 structure and the experimentally-derived structure, we find that the rho was not impacted by the difference in the distance per residue for up to 4Å (S2B Fig). There is, however, a positional effect on the heterogeneity, with all amino acids at specific positions contributing a vast portion of the heterogeneity in the prediction from FoldX and Rosetta.

### Comparison of predicted |ΔΔG| from experimentally-derived structures vs AF2 structures to predict LOF

The predicted |ΔΔG| from FoldX analyzed with the AF2 structures were significantly better than the experimentally-derived structures in the BRCT domain of *BRCA1* and the DNA binding domain (DBD) binding domain in BRCA2 to predict the categorical LOF activity using the De-long test to compare the AUC (Fig 2A). In the BRCT domain of *BRCA1*, FoldX generated an AUC=0.861 (95% CI:0.858-0.863) from the predicted AF2 complex structure of 4OFB, compared to AUC=0.847 (95% CI: 0.844-0.850) from the experimentally-derived complex structure of 4OFB. FoldX also generated an AUC=0.836 (95% CI:0.833-0.839) from the AF2 subunit structure of IJNX, compared to an AUC=0.792 (95% CI:0.789-0.795) from the experimentally-derived subunit structure of 1JNX, showing that utilizing the complex AF2 structure enhances the prediction of LOF activity using FoldX in the BRCT domain of *BRCA1*. However, in the RING domain of *BRCA1*, FoldX using complex experimentally-derived structure of 7LYB was significantly better than the predictions from FoldX with an AF2 wildtype template of 7LYB, with an AUC=0.835 (95% CI=0.831-0.840), compared to AUC=0.741 (95% CI:0.735-0.748). The AF2 complex structure of 7LYB was noted to have a mean per residue distance of 19Å when superimposed on the experimentally-derived structure, which was outlier compared to the other 6 structures that had a mean per residue distance of 3Å (S3 Fig).

**Fig 2:**
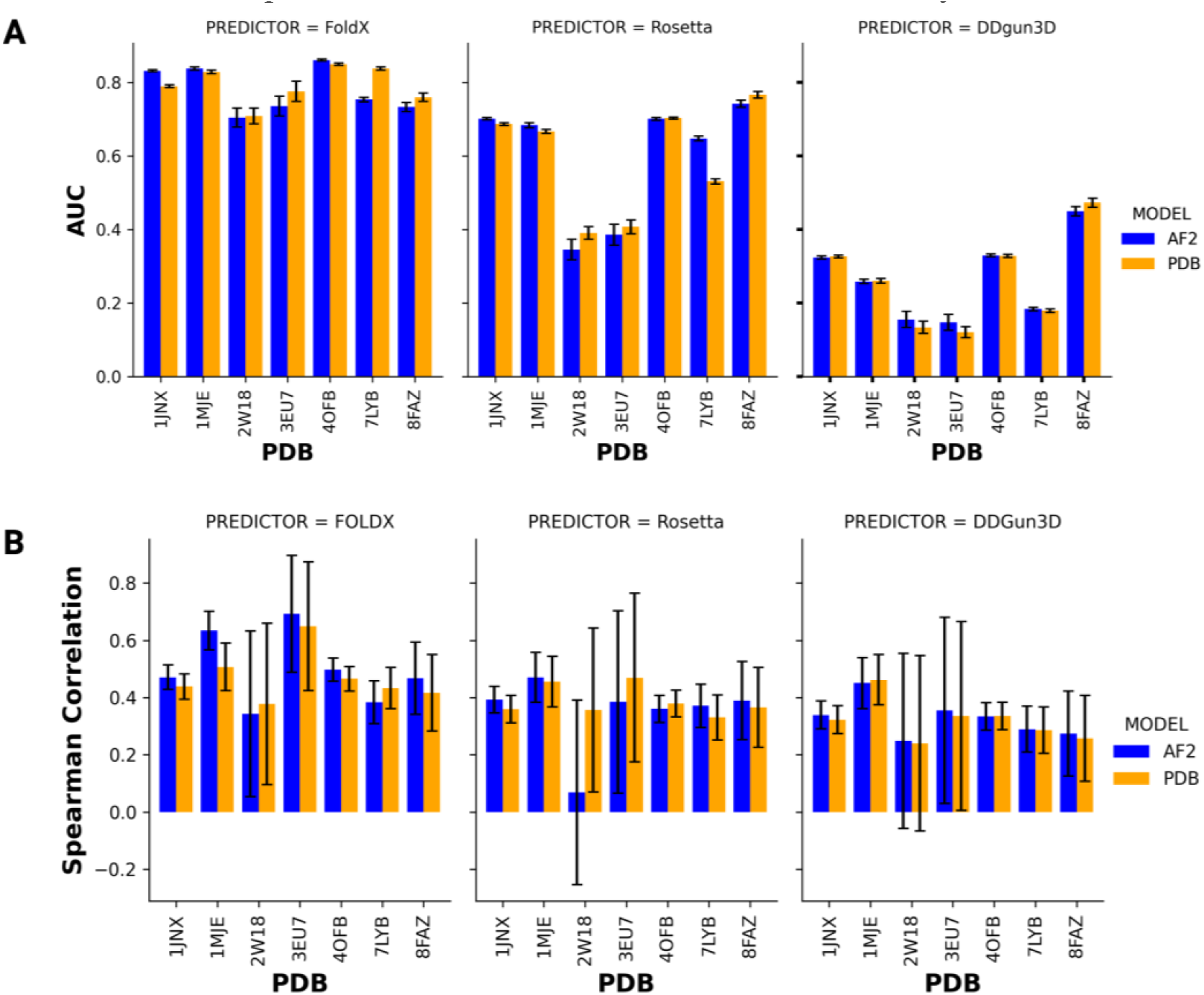
|ΔΔG| prediction of Loss-of-function activity on a categorical and continuous scale. (A) Area under curve denoting the predictive ability of |ΔΔG| from FoldX, Rosetta and DDGun3D using Experimentally-derived structures vs AF2 structures as the wild-type template to predict loss-of-function activity in *BRCA1, BRCA2, PALB2* and *RAD51C*. (B)Spearman correlation of predicted |ΔΔG| from FoldX, Rosetta and DDGun3D using Experimentally-derived structures vs AF2 structures with functional HDR measurements in *BRCA1, BRCA2, PALB2* and *RAD51C*.

In *BRCA2*, Rosetta analyzed with the AF2 complex structure of 1MJE representing the BRCA2-DSS1-ssDNA complex, generated an AUC=0.681 (95% CI:0.675-0.687), which was significantly better that an AUC=0.665 (95% CI:0.659-0.672) which was generated when analyzed by the experimentally-derived complex structure of 1MJE. FoldX predictions of LOF activity were significantly better than Rosetta and DDGun3D, and sub-setting to the mutations to the DNA binding domain (DBD) of BRA2, FoldX analyzed with AF2 complex structure of 1MJE generated an AUC=0.8364 (95% CI:0.8322-0.8407), which was significantly better than an AUC=0.8299 (95% CI:0.8213-0.8320) that was generated by FoldX analyzed with the experimental-derived complex structure of 1MJE.

In *PALB2*, there was no significant difference between the predicted |ΔΔG| from the AF2 complex structure and the experimentally-derived complex structure analyzed with FoldX. Among the stability predictors, FoldX predictions of LOF activity were significantly better than Rosetta and DDGun3D. Analyzing the AF2 complex structure of 3EU7, FoldX generated an AUC=0.721 (95% CI:0.694-0.747), compared to Rosetta and DDGun3D, which generated an AUC=0.401 (95% CI:0.375-0.427) and AUC=0.157 (95% CI:0.135-0.179) respectively. With the AF2 subunit structure of 2W18, FoldX generated an AUC=0.719 (95% CI: 0.695-0.743), which was significantly better than Rosetta and DDgun3D, with an AUC=0.359 (95% 0.329-0.388) and AUC=0.159 (95% CI:0.138-0.181) respectively.

In *RAD51C*, we find that predicted |ΔΔG| analyzed from the experimentally-derived complex structure of 8FAZ was significantly better than the complex AF2 structure of 8FAZ analyzed with Rosetta and FoldX to predict LOF activity. An AUC=0.766 (95% CI:0.757-0.775) was generated from experimentally-derived complex, which was significantly better than AUC=0.741 (95% CI:0.731-0.751) generated from the AF2 structure analyzed with Rosetta. With FoldX as well, an AUC=0.759 (95% CI: 0.748-0.771) was generated from experimentally-derived complex, which was significantly better than AUC=0.732 (95% CI: 0.720-0.745) from the AF2 complex.

Using spearman correlation coefficient (rho) as the metric, we evaluated the prediction of |ΔΔG| using the experimentally-derived structures and AF2 structures with the continuous measurement of HDR functional activity from MAVE assays and found no significant difference in the correlation of functional activity with |ΔΔG| from the 7 AF2 structures or its experimentally derived counterparts analyzed with FoldX, Rosetta or DDGun3D (Fig 2B). However, there was heterogeneity from the prediction among the stability predictors. The rho values derived using the predicted |ΔΔG| analyzed from the AF2 complex structure of 4OFB representing *BRCA1* BRCT domain and AF2 structure of 1MJE representing BRCA2-DSS1-ssDNA complex using FoldX was significantly better than the rho from the AF2 structures or the experimentally-derived structure analyzed with Rosetta and DDGun3D. A rho=0.497 (95% CI:0.455-0.537) was observed from the AF2 structure of 4OFB, compared to rho=0.379 (95% CI:0.331-0.424) from experimentally-derived structure and rho=0.360 (95% CI:0.312-0.407) from the AF2 structure analyzed by Rosetta, and rho=0.33 (95% CI:0.25-0.382) from experimentally-derived structure and rho=0.334 (95% CI:0.285-0.381) from the AF2 structure analyzed by DDGun3D. A rho=0.634 (95% CI:0.562-0.696) was observed from the AF2 structure of 1MJE, compared to rho=0.455 (95% CI:0.362-0.539) from experimentally-derived structure and rho=0.470 (95% CI:0.378-0.553) from the AF2 structure analyzed by Rosetta, and rho=0.462 (95% CI:0.369-0.545) from experimentally-derived structure and rho=0.470 (95% CI:0.357-0.535) from the AF2 structure analyzed by DDGun3D. In *PALB2*, and *RAD51C*, there was no significant difference between FoldX, Rosetta and DDGun3D to predict the continuous HDR functional activity.

### Comparison of protein stability predictors with *in silico* missense predictors

We find that existing *in silico* missense predictors significantly predict LOF better than protein stability predictors based on the area under the curve (AUC) (Fig 3A). We also find that no single *in silico* missense predictor predicts LOF the best based on the AUC for all 4 genes. Using the average AUC from all 4 genes as the metric, we find that MetaRNN and AlphaMissense predict LOF activity the best, with MetaRNN having an average AUC=0.891 (95% CI:0.886-0.895) and AlphaMissense having an average AUC=0.890 (95% CI 0.886-0.895) across all 4 genes. To conclude which is the best *in silico* missense or stability predictor to use to predict LOF on across all 4 genes, we rank ordered the predictions for all predictors in the dbNSFP database and stability predictors and calculated the average rank across all 4 genes. From our results (Fig 3B) we find that AlphaMissense had the highest average rank, followed by MetaRNN, and BayesDel_addAF. This result would imply that AlphaMissense is the best *in silico* missense predictor in predicting LOF activity in these breast cancer genes. The best stability predictor is FoldX using AF2 wild-type structures which had a rank of 19. This suggests that predicted |ΔΔG| predicts LOF better than 38 other predictors in the dbNSFP database. However, we should note that the mutations used for this evaluation are enriched with mutations that are in ordered regions of the protein.

**Fig 3:**
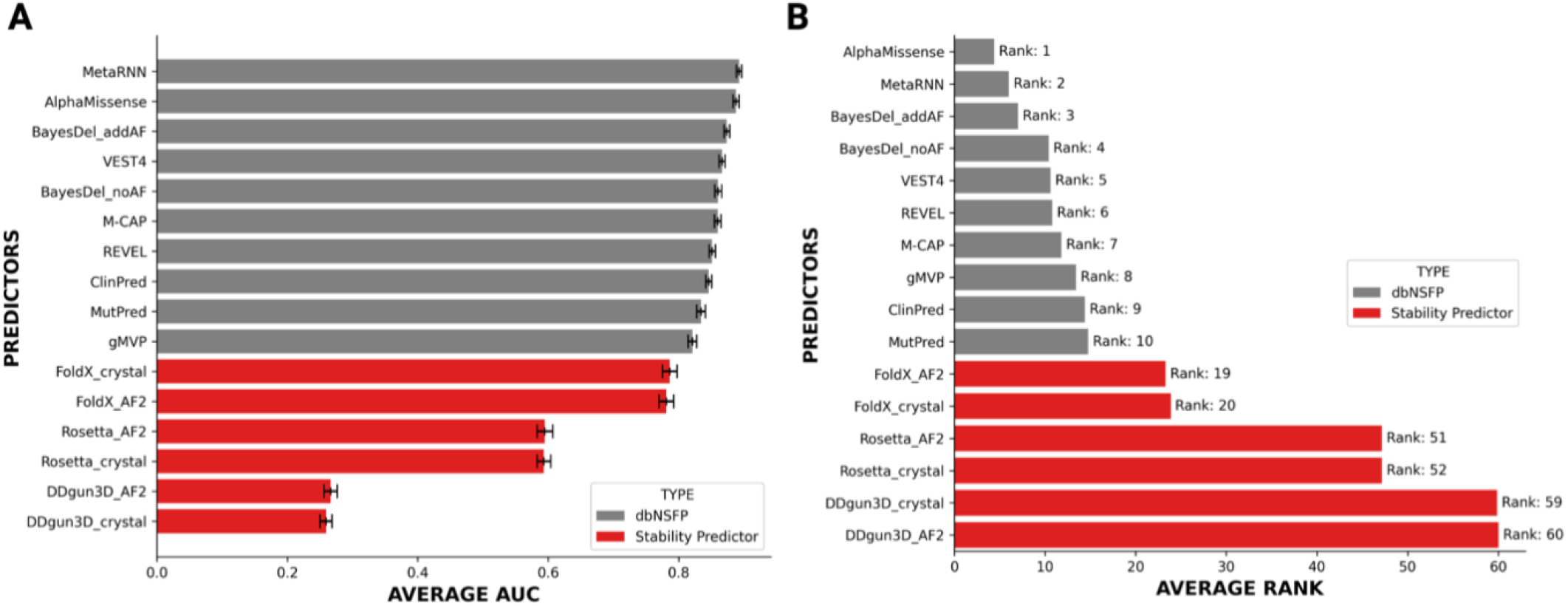
Performance of *in silico* predictors to measure loss-of-function activity in *BRCA1, BRCA2, RAD51C* and *PALB2*. (A): Area under the curve of *in silico* missense predictors vs stability predictors to predict loss-of-function activity in *BRCA1, BRCA2, RAD51C* and *PALB2*. (B) Rank ordered performance of *in silico* missense predictors vs stability predictors to predict loss-of-function activity in *BRCA1, BRCA2, RAD51C* and *PALB2*.

## Discussion

### AF2 predictions are better than ESMFold

AF2 and ESMFold are considered to two best protein structural predictors and consistent with existing literature, AF2 multimer predicts protein structures that are highly similar to experimentally-derived structures compared to ESMFold in *BRCA1, BRCA2, PALB*2, and *RAD51C*. ESMFold predictions are comparatively faster than AF2, but the predictions are limited to 1024 amino acids, thereby larger structures, such as 7LYB were not compared. AF2 is more suited for this study for the high similarity and its ability to predict large protein structures.

### Association of predicted protein stability to protein Loss-of-function

Consistent with existing literature, we found that there is a strong association of stability towards LOF activity using all three protein stability predictors, suggesting that protein destabilization or over-stabilization is an important factor in the contribution to LOF activity in *BRCA1, BRCA2, PALB2* and *RAD51C*[7, 8]. These genes have been reported to be haploinsufficient, suggesting that minor perturbation is required to completely destabilize the protein to impact loss-of-function. We also chose FoldX, Rosetta and DDGun3D as these predictors were shown to perform the best in terms of correlation results with deep mutational scan data using complex experimentally-derived protein structures and these predictors require a wildtype protein template to generate its predictions[16].

### Prediction from DDGun3D is less dependent on wild-type protein templates

Overall, strong correlation was observed between the |ΔΔG| prediction using either AF2 structure of the experimentally-derived structure, with the exception for 7LYB and 1MJE. We find that DDGun3D predictions showed stronger correlations compared to FoldX and Rosetta. This could be due to the fact that only 33% of DDGun3D predictions is based on the wildtype protein structure, while the remaining contribution comes from Blosum62 matrix substitution scores, difference in statistical potential score from the linear chain of the amino acid between the wild-type and mutant, and the difference between the hydrophobicity of the wild-type and mutant[15], whereas FoldX and Rosetta use a mixture of physics and statistical methods to compute protein stability based on the clashes caused by introducing a mutant within the wild-type structures.

### Protein structural features have no impact on protein stability predictions

Recent publications have highlighted the use of protein structural features from AF2 structures to study LOF activity[25]. Features such as PLDDT, RSA, distance between residue, physicochemical properties such as atomic weight, isoelectric points and aromaticity were shown to be useful in predicting LOF activity. We tested the dependence of the difference in |ΔΔG| prediction from experimentally-derived structure and AF2 structures with these features. Our results showed no influence on the difference in stability prediction to the difference in features values, thus suggesting that these features are independent of protein stability and hence have independent effects on the relationship with LOF. Thereby we can make a claim that stability, along with features such as PLDDT, RSA, distance between residue, physicochemical properties such as atomic weight, isoelectric points and aromaticity can be used to study protein LOF in *BRCA1, BRCA2, PALB2* and *RAD51C*.

### FoldX prediction using AF2 wild type structures predicts loss-of-function better than other stability predictors

Overall, our results support the choice of using FoldX as the protein stability predictor in these genes using complex structures of AF2 predicted structures, compared to Rosetta and DDGun3D. More importantly, our results suggest that using FoldX with AF2 structures that are highly similar to experimentally-derived structures predicts LOF activity just as good as prediction from FoldX with complex experimentally-derived structures. Our results also suggest that there is a domain specific effect on the predictiveness of stability towards function in *BRCA1*, suggesting that the degree of perturbation is heterogeneous across domains. Even though the predictions are based on several replicates, we acknowledge that there is inherent noise in the predictions from FoldX, Rosetta and DDGun3d. These predictors were not inherently built to predict protein LOF, but experimental protein stability.

### AlphaMissense is the best *in silico* predictor in breast cancer genes

*In silico* missense predictors predict function better than protein stability predictors in *BRCA1, BRCA2, RAD51C* and *PALB2*. AlphaMissense ranks highly in the prediction of LOF activity in all 4 genes and on average AlphaMissense ranks the best. BayesDel is currently considered the *in silico* model of preference by the ClinGen *BRCA1/2* VCEP (https://cspec.genome.network/cspec/ui/svi/doc/GN092), however newer predictors AlphaMissense and MetaRNN predict function significantly better than BayesDel in these predisposed breast cancer genes. AlphaMissense utilizes a transformer based multiple sequence alignment of protein sequences, whereas MetaRNN is an ensemble neural network model that uses many existing *in silico* missense predictors as features and hence there is an argument that the predictions might be overfit to the training data to the individual features.

In summary, Generative AI tools such as AF2 multimer predictions can be used in the prediction of protein stability in haploinsufficient genes *BRCA1, BRCA2, PALB2* and *RAD51C* to study LOF activity. However, these predictions are primarily focused on mutations in ordered regions and do not account for disordered regions of the protein complex. The application of generative AI to study functions in disordered regions was beyond the scope of this study. Additionally, Multiplexed Assays for Variant Effects (MAVE) provide an independent dataset for evaluating LOF activity, though these assays are known to have errors due to the type of assay used, experimental artifacts, and variability in replicates. While these assays will not replace computational methods in the near future, they contribute to refining our understanding of the factors causing deleterious effects on protein function.

## Methods

### Data Selection

We used the functional classification derived from the MAVE assays in genes *BRCA1, BRCA2, PALB2* and *RAD51C* respectively and the classification from the ClinVar database (S1 Table). The MAVE assay for *BRCA1* was based on saturation Genome editing in HAP1 cell lines that had a total of 2086 mutations over the RING and BRCT functional domains that aim to measure Loss-of-function in this cell line [27]. For *BRCA2*, we used the functional classification from a homology-directed repair (HDR) cell-based assay from 462 missense variants affecting the *BRCA2* DNA binding domain [28]. For *RAD51C*, we used the functional classification from a homology-directed repair (HDR) reporter assay, which introduced 174 missense variants in mammalian hRAD51C expression constructs using site-directed mutagenesis [29]. For *PALB2*, we used the functional classifications from 91 missense variants evaluated from a Homology directed repair (HDR) assay[30, 31].

We further supplemented our dataset with the mutational classification from the ClinVar database listed in dbNSFP 202403 release [45]. We group all classification of pathogenic and likely pathogenic into the “deleterious” category, and classification of benign and Likely benign into the “neutral” category.

### PDB Selection

We chose 8 structures from the PDB (S2 Table), that represent different ordered regions of genes *BRCA1, BRCA2, RAD51C* and *PALB2*. We chose these structures based on the resolution and coverage. These structures exist as a subunit and as a complex, with two or more protein chains representing a complex or as a single chain representing a subunit. For *BRCA1*, we downloaded 4OFB, 1JNX, and 7LYB, as these structures represent the two ordered domains of *BRCA1* (BRCT and RING). 1JNX is a single chain subunit structure and 4OFB is a complex that contains 2 protein chains, both representing the BRCT domain of *BRCA1*. 7LYB is a 7-protein chain complex structure, which chain ‘M’ representing the RING domain of *BRCA1*. For *BRCA2*, we downloaded 1MJE, which is a two-protein chain complex structure, and chain “C” represents the ordered regions of the DBD domain of *BRCA2*. For *RAD51C*, we downloaded 8FAZ, a complex structure that contains 4 protein chains and with chain “C” representing the entire gene of RAD51C. For *PALB2*, we downloaded 3EU7, a complex structure with two chains, with chain “A” representing the entire *PALB2* gene. We also downloaded 2W18, a subunit structure representing the *PALB2* gene.

### AlphaFold2 and ESMFold prediction

AF2 structures of the 8 high resolution PDB structures were generated using ColabFold v1.5.5 [46] using the pdb100 templates. Default settings were used for the remaining configurations. Using BioPython[47], the FASTA sequence from each experimentally-derived structure was extracted and used as inputs to generate five separate replicates, ranked based on model confidence. For the eight experimentally-derived structures 8FAZ, 8FOUY, 7LYB, 4OFB, 1JM7, 1IYJ, 1MJE, 3EU7, and 2W18, we generated predicted protein structures using AF2 multimer predictions with ColabFold.

Using the Python implementation of ESMFold, we generated predicted structures for six out of the eight experimentally-derived structures using their FASTA sequences as input. We employed the ESM-2 pretrained model “esm2_t33_650M_UR50D,” which comprises 650 million parameters and 33 layers, running on a single T4 GPU. ESMFold has a limitation of generating structures with a maximum of 1024 amino acids. Consequently, for the complex structures 7LYB and 1IYJ, which exceed 1024 amino acids, complete structures were not created. Full details can be found in the code found in https://github.com/rohandavidg/CarePred.

### Protein Stability

FoldX, Rosetta and DDGun3D predictors were used to predict ΔΔG with both AF2 and experimentally-derived structures for all possible missense mutations, leading to a total of 246,910 mutations. Using the python module pyFoldX [48], which uses FoldX 5.0 the structures were passed through the “RepairPDB ‘‘ function prior to ΔΔG calculations, with default settings. For every mutation, five replicates were computed and the mean ΔΔG value was calculated. DDGun3D python package was used to generate predicted ΔΔG using the default settings, five replicates were computes and the mean ΔΔG value was calculated. The Cartesian ΔΔG application from Rosetta suite (Linux build 2021.16.61629) was used to generate the ΔΔG predictions, following standard protocols published in several publications [49, 50] while using the Ref2015 scoring function. The structures were initially relaxed and ΔΔG predictions were made over three iterations, which was averaged to generate mean ΔΔG in (Rosetta energy per unit). To convert the units into kcal/mole scale, we used a scaling factor of 2.94, as shown in the literature [16, 49].

### Statistical analysis

A Mann-Whitney U statistical test was employed from the SciPy[51] stats python package to understand the association of predicted |ΔΔG| from FoldX, Rosetta and DDGun3D to protein LOF activity. The metric RMSD calculations were computed using the *superimposer* method from the BioPython PDB python package. RMSD was used as the metric to evaluate the similarity between the AF2 structure compared to the experimentally-derived structure downloaded from the PDB. Similarity was calculated over the entire structure and over the Cα backbone chain. We use SciPy stats package to compute the spearman correlation coefficient between predicted |ΔΔG| from AF2 structures and experimentally-derived structures analyzed with FoldX, Rosetta and DDGun3d to evaluate the linearity. We also compute the spearman correlation of ΔΔG from AF2 structures and experimentally-derived structures analyzed with FoldX, Rosetta and DDGun3d, compared to the continuous functional MAVE activity score. To compute the confidence interval of the spearman correlation, we do a Fischer transformation of these correlation coefficients. Utilizing these Fisher transformations, we calculate the combined standard errors, the z-scores of the difference and then using the z-scores, we calculate the p-value based on the normal distribution.

To calculate the predictive ability of missense *in silico* predictors and stability predictors to LOF activity, we use the metric AUC from the *roc_auc_score* method from the scikit-learn [52] package. We compute the 95% confidence intervals by sampling 1000 times across the dataset.

## Supplemental

**S1 Fig:**
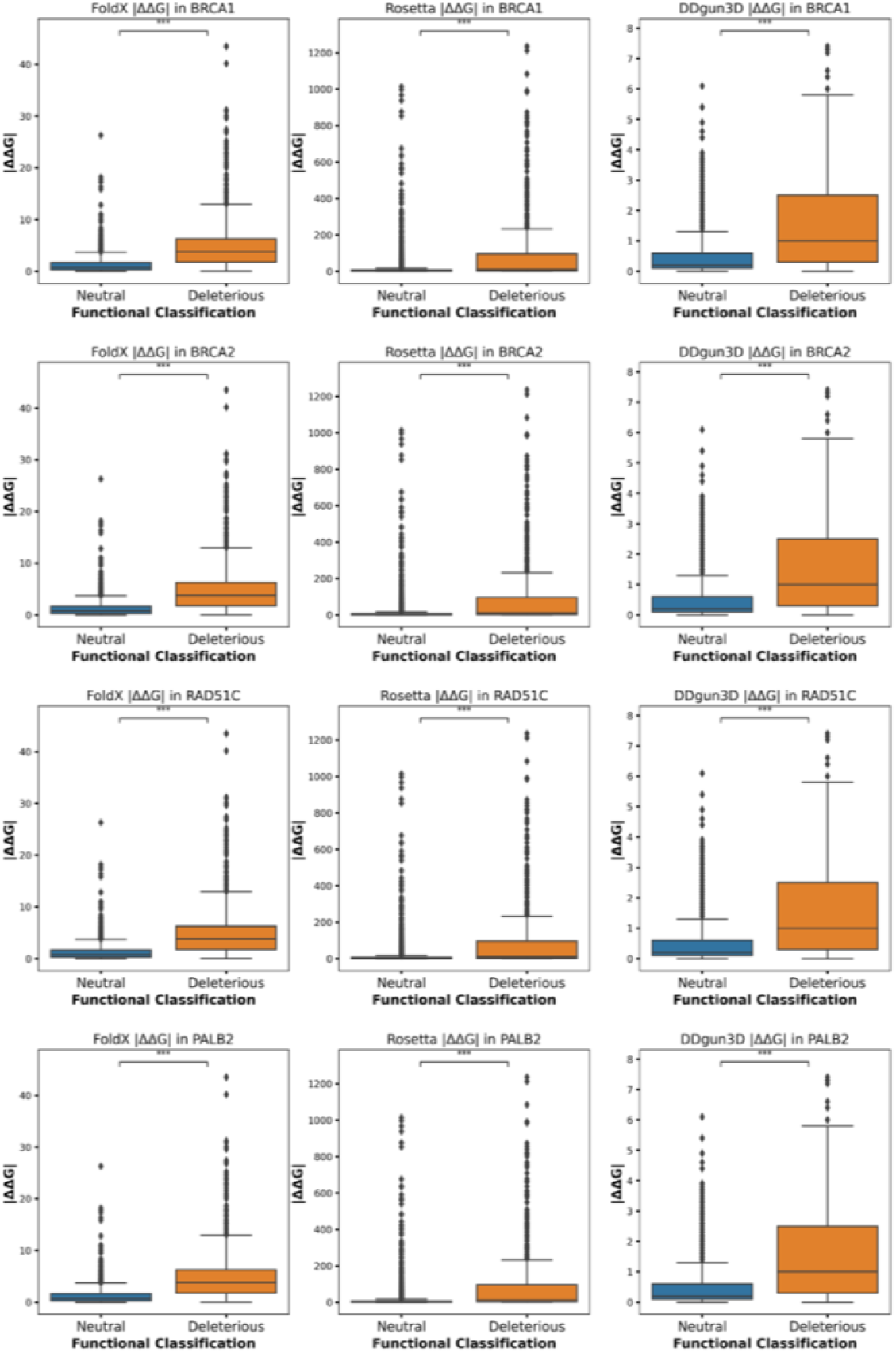
Distribution of predicted |ΔΔG| from FoldX, Rosetta and DDGun3D stratified by functional classification (Deleterious vs Neutral) in genes *BRCA1, BRCA2, PALB2* and *RAD51C*, with the association between the two groups denoted by the Mann-Whitney U test.

**S2 Fig:**
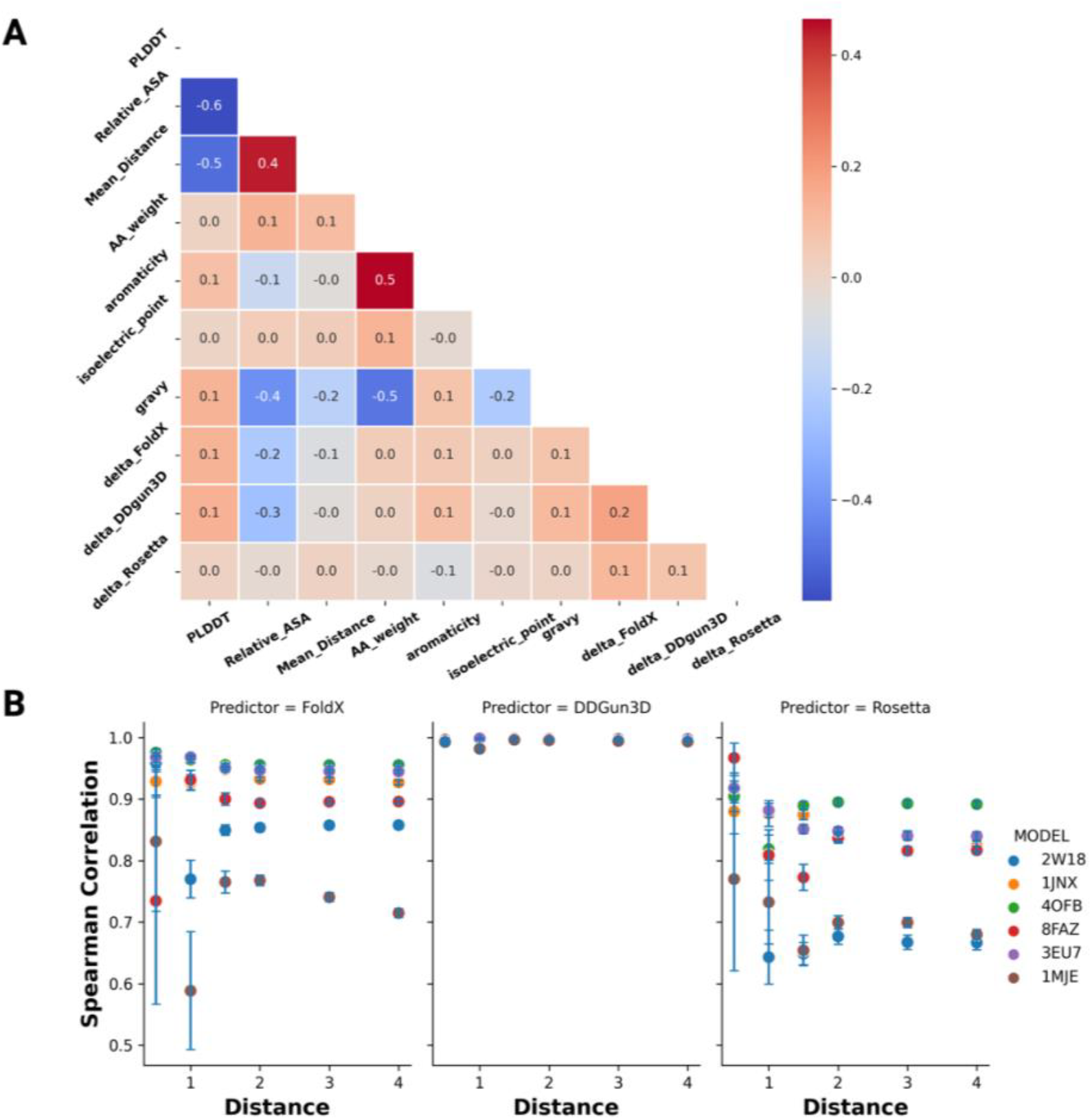
Linear association of difference between the ΔΔG from AF2 structures vs experimentally-derived structure as the wild-type template analyzed with FoldX, Rosetta and DDGun3D with the features extracted from AF2 structure and per residue distance between the superimposed AF2 structure onto the experimentally-derived structure. (A). Spearman correlation heatmap of feature derived from AF2 structures and deltas of ΔΔG derived from AF2 structures and experimentally-derived structures analyzed with FoldX, Rosetta and DDGun3D (B) Spearman correlation scatterplot of the difference between the ΔΔG from AF2 structures vs experimentally-derived structure as the wild-type template analyzed by FoldX, DDGun3D and Rosetta stratified by the per residue distance when the AF2 and experimentally-derived structure were superimposed on each other.

**S3 Fig:**
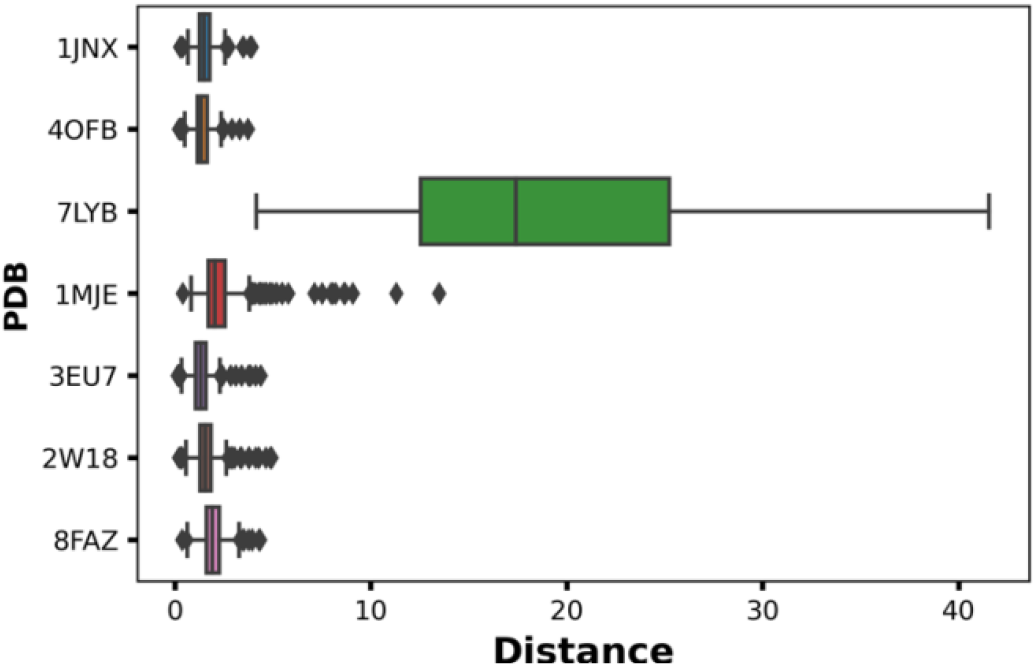
Distribution of the per residue distance in Å between AF2 predicted structure and experimentally-derived structure found in the PDB. AF2 prediction of 7LYB was an outlier with a mean distance of 19Å

**S1 Table:**
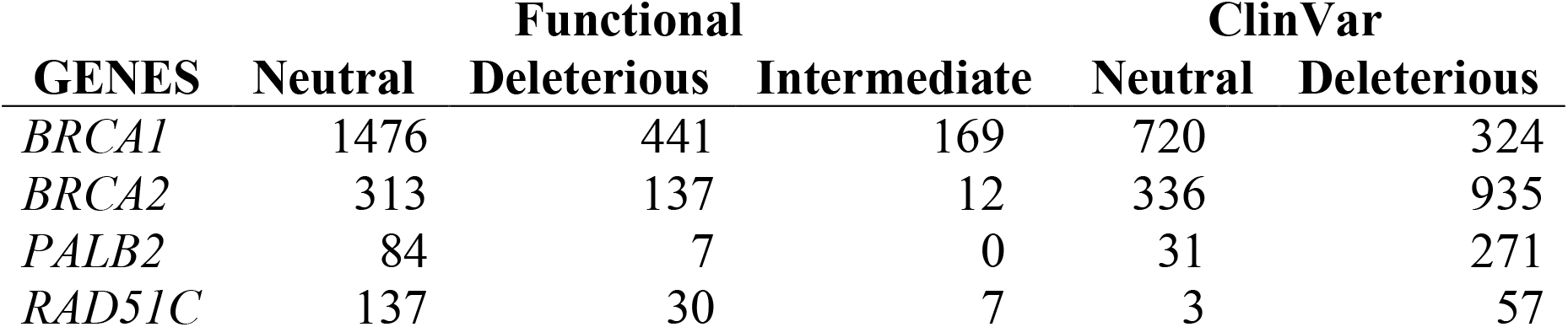
Table denoting the total number of classified mutations used from the ClinVar database and MAVE functional assays for evaluation.

**S2 Table:** List of PDB ID from Genes used in this study.

| Gene | PDB ID | Atom count | Resolution | Residue count | Protein chains |
| --- | --- | --- | --- | --- | --- |
| *RAD51C* | 8FAZ | 7331 | 2.30 Å | 926 | 4 |
| *BRCA1* | 7LYB | 14699 | 3.28 Å | 1389 | 7 |
| *BRCA1* | 4OFB | 1791 | 3.05 Å | 223 | 2 |
| *BRCA1* | 1JNX | 1660 | 2.50 Å | 207 | 1 |
| *BRCA2* | 1IYJ | 10092 | 3.40 Å | 1272 | 2 |
| *BRCA2* | 1MJE | 5198 | 3.50 Å | 648 | 2 |
| *PALB2* | 3EU7 | 2585 | 2.20 Å | 327 | 2 |
| *PALB2* | 2W18 | 2543 | 1.90 Å | 306 | 1 |

